# Human ESCRT-III Polymers Assemble on Positively Curved Membranes and Induce Helical Membrane Tube Formation

**DOI:** 10.1101/847319

**Authors:** Aurélie Bertin, Nicola de Franceschi, Eugenio de la Mora, Sourav Maity, Nolwen Miguet, Aurélie di Cicco, Wouter Roos, Stéphanie Mangenot, Winfried Weissenhorn, Patricia Bassereau

## Abstract

Endosomal sorting complexes required for transport-III (ESCRT-III) are thought to assemble *in vivo* inside membrane structures with a negative Gaussian curvature. How membrane shape influences ESCRT-III polymerization and conversely how ESCRT-III polymers shape membranes is still unclear. Here, we used human core ESCRT-III proteins, CHMP4B, CHMP2A, CHMP2B and CHMP3 to address this issue *in vitro* by combining membrane nanotube pulling experiments, cryo-electron microscopy, cryo-electron tomography and high-speed AFM. We show that CHMP4B filaments bind preferentially to flat membranes or to membrane tubes with a positive mean curvature. Both CHMP2B and CHMP2A/CHMP3 assemble on positively curved membrane tubes, the latter winding around the tubes. Although combinations of CHMP4B/CHMP2B and CHMP4B/CHMP2A/CHMP3 are recruited to the neck of pulled membrane tubes, they also reshape large unilamellar vesicles into helical membrane tubes with a pipe surface shape. Sub-tomogram averaging reveals that the filaments assemble parallel to the tube axis with some local perpendicular connections, highlighting the particular mechanical stresses imposed by ESCRT-III to stabilize the corkscrew-like membrane architecture. Our results thus underline the versatile membrane remodeling activity of ESCRT-III that may be a general feature of ESCRT-III required for all or selected cellular membrane remodeling processes.

## Introduction

The Endosomal Sorting Complex Required for Transport (ESCRT-III) is an ancient conserved membrane remodeling machine. ESCRT-III employs polymer formation to catalyze inside-out membrane fission processes in a large variety of cellular processes, including budding of endosomal vesicles and enveloped viruses, cytokinesis, nuclear envelope reformation, plasma membrane and endolysosomal repair, exosome formation, neuron pruning, dendritic spine maintenance and preperoxisomal vesicle biogenesis ^1–12^. Yeast ESCRT-III comprises four subunits Vps20, Snf7, Vps2 and Vps24, which polymerize in this order on endosomal membranes ^13^, and is dynamically regulated by the ATPase VPS4 ^14^. The corresponding human homologues comprise one or several isoforms named CHMP6 (Vps20), CHMP4A, B, C (Snf7), CHMP2A, B (Vps2) and CHMP3 (Vps24) in addition to CHMP1A, B, CHMP5, CHMP7 and CHMP8/IST1 ^6^. ESCRT-III proteins adopt an auto-inhibited conformation in the cytosol ^15–17^, which requires the release of the C-terminal auto-inhibition ^18, 19^. This leads to polymerization of loose CHMP4 spirals ^20–22^, helical CHMP2A-CHMP3 spirals ^16, 23, 24^, CHMP2A filaments ^25^ and Vps24 (CHMP3) filaments ^26^ *in vitro*. *In vivo*, CHMP4 or CHMP2B over-expression leads to membrane tube formation with CHMP4 and CHMP2B filaments inside the tube, respectively ^27–29^. Polymerization is guided by conformational changes that stabilize the filaments via domain exchange, thereby generating basic surfaces for interaction with positively curved ^30^ or negatively curved membranes ^31^ carrying a negative net charge ^30, 32, 33^. Although Snf7 (CHMP4) polymerizes on supported lipid bilayers ^22^, preformed membrane curvature was suggested to favor Snf7 (CHMP4) membrane interaction ^34, 35^.

Common to all ESCRT-mediated processes is the strict requirement of VPS4 that not only recycles ESCRT-III ^36^, but actively remodels the polymers *in vivo* ^14, 37^ and *in vitro* ^38, 39^. Vps4/VPS4 ESCRT-III remodeling seems to be critical to complete release ^14, 40, 41^. Furthermore, all ESCRT-catalyzed processes recruit CHMP4 and downstream CHMP2 isoform(s) ^3^, indicating CHMP4 and CHMP2 are core components for ESCRT-III function. Accordingly, HIV-1 budding requires only one CHMP4 and CHMP2 isoform for virus release ^42, 43^, although other ESCRT-III members are recruited during HIV-1 budding ^44–46^ and enhance budding efficiency ^25^. This thus suggests a minimal budding/membrane fission machinery that requires CHMP4 and CHMP2 isoforms. Consistent with this proposal, *in vitro* reconstitution experiments implicated Snf7 (CHMP4), Vps24 (CHMP3) and Vps2 (CHMP2A) in the Vps4-driven release of membrane tubes *in vitro* ^33^. Based on the core fission machinery a number of different models have been proposed to explain ESCRT-catalyzed membrane fission ^6, 10, 47^.

*In vivo* filament assembly has been imaged within bud necks of viruses ^43, 45, 48^. Similarly, ESCRT-III containing spirals have been observed within the cytokinetic midbody ^49–52^, and proposed to be multi-stranded ^37, 49^. This thus suggests that ESCRT-III assembles on membranes that exhibit a saddle-like shape with negative Gaussian curvatures (See Supplementary Fig. S1 for a definition of the different curvatures involved in this work).

Although the intrinsic curvature of the filaments and their flexibility are likely important to shape membranes, the precise role of ESCRT-III polymers and their preference for different or the same membrane geometries is still unclear ^22, 24, 34, 53^.

Here we investigated how ESCRT-III polymerization shapes membranes and how membrane shape influences their assembly on membranes. To address these questions, we have developed *in vitro* assays based on the essential core of purified human ESCRT-III proteins (CHMP4B, CHMP2A, CHMP2B and CHMP3) and model membrane systems. We have used C-terminally truncated versions of CHMP2A and CHMP2B to facilitate polymerization as well as full-length CHMP4B and CHMP3. We have designed confocal microscopy experiments with membrane nanotubes of controlled geometries pulled from Giant Unilamellar Vesicles (GUVs) to study the effect of membrane mean curvature and topology on ESCRT-III protein recruitment and polymerization at the macroscopic scale. Furthermore, by using high-speed AFM (HS-AFM) and cryo-electron microscopy (cryoEM), we have obtained nanometer resolution images showing the preferential membrane shape induced upon ESCRT-III assembly on small liposomes and preformed tubes and the corresponding organization of the protein filaments at their surface.

## Results

### CHMP4B polymers do not deform membranes and have no particular preference for curved membranes

First, we have confirmed with high-speed AFM that the human CHMP4B, similarly to the yeast Snf7, assembles into spirals when in contact with a negatively-charged supported lipid bilayer (SLB) made of 60% DOPC, 30% DOPS, and 10% PI(4,5)P2 (Fig. 1A and Supplementary Movie S1). We measured an average peak to peak distance between filaments within a spiral equal to 11.3 ± 1.9 nm (N = 134) (Supplementary Fig. S2), slightly smaller than reported for Snf7 (17 ± 3 nm) ^22^. Thus, CHMP4B spirals are built up from one single, unbranched filament, forming a tighter structure than Snf7 spirals that display inter-filament branching ^22^. In order to test if CHMP4 can deform membranes as a “loaded spring” as previously suggested ^54^, we have employed an *in vitro* assay involving deformable vesicles. We have analyzed both the membrane deformation and the organization of CHMP4B filaments by cryoEM, at nanometer scales. LUVs (Large Unilamellar Vesicles of 50 nm to 1 µm in diameter) made of 70 % EPC, 10 % DOPE, 10 % DOPS, 10% PI(4,5)P2 were incubated with CHMP4B, plunged frozen and imaged by cryo-EM (N = 8 experiments). As displayed in Fig. 1B, CHMP4B assembles into spirals on LUVs. The inter-filament distances within the spirals equals to 7.8 ± 2.6 nm (N = 208), corresponding to peak-to-peak distances of 11.3 ± 2.6 nm, similar to those measured on rigid substrates with HS-AFM. The diameter of the spirals is approximately 193 ± 63 nm (N = 23). We did not observe any obvious membrane budding or buckling but rather an apparent flattening as compared to control LUVs (Supplementary Figs. S3A-B). To unambiguously visualize any 3D deformation, we have performed cryo-electron tomography (cryoET) (Fig. 1C and Supplementary Movie S2) (N = 5). When bound to membranes, CHMP4B spirals (in red) are clearly flat and follow the contour of vesicles without inducing any noticeable deformation as highlighted in the side view (Fig. 1C, bottom) and in Supplementary Movie S2. This suggests that the elastic energy stored in the spirals favors a non-curved membrane and no invagination or ex-vagination.

**Figure 1:**
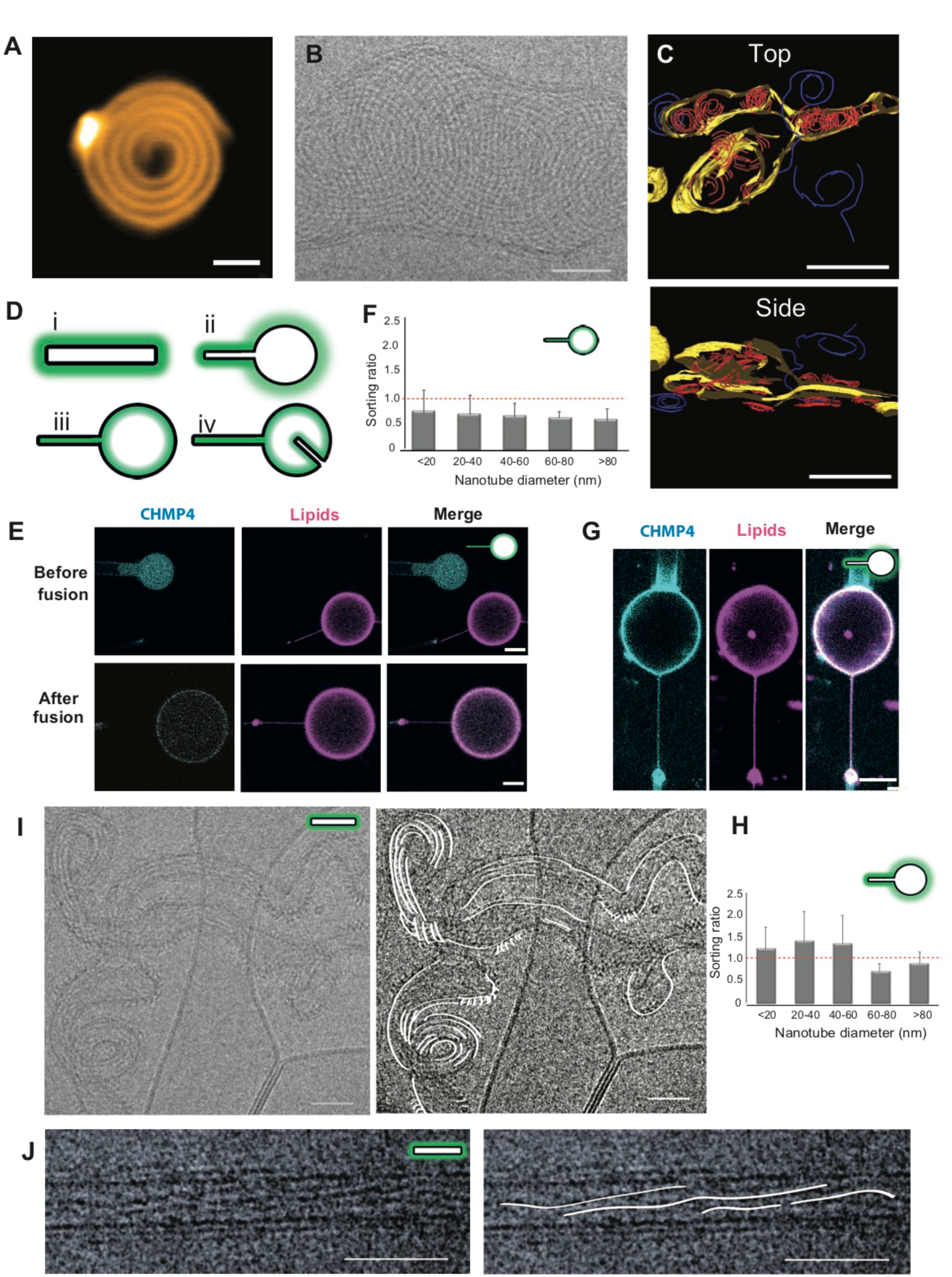
CHMP4 flattens LUVs and binds preferentially to flat membranes or to membranes with a positive mean curvature. **A**: CHMP4B spirals observed by HS-AFM on a supported lipid bilayer. Scale bar: 50 nm. **B**: Cryo-EM image of CHMP4B spiral polymers on deformable LUVs. Scale bar: 50 nm. **C**: Top view (top) and side view (bottom) of a cryo-EM tomogram showing CHMP4B spirals on deformable LUVs (red: CHMP4 filaments polymerized on lipids; blue: filaments polymerized on the grid; yellow: lipids) (corresponds to Supplementary Movie S2). Scale bar: 200 nm. **D**: Schematic depicting the different geometries used to study binding and polymerization of the ESCRT-III proteins. In each case, the location of the protein inside or outside the GUV is indicated by a green shadow. (i) Proteins outside a free isolated tube, corresponding to C>0 and K=0. (ii) Proteins outside a nanotube pulled from a GUV: on the tube, C>0 and K=0; on the GUV, C= K=0; and on the neck, K<0. (iii) Proteins inside a nanotube pulled from a GUV: on the tube, C<0 and K=0; on the GUV, C= K=0; and on the neck, K<0. (iv) Spontaneously formed tubule inside a GUV in geometry (iii): on the internal tube, C>0 and K=0. **E**: Confocal images corresponding to a GUV fusion experiment in which CHMP4B is binding in geometry (iii). Scale bar: 10µm. **F**: Quantification of the sorting ratio for 17 nanotubes from 17 GUVs in 8 independent GUV preparations and variable diameters, corresponding to the experiment type shown in Figure 1E. For each nanotube diameter, N measurements have been obtained: <20nm: N=19; 20nm-40nm: N=28; 40nm-60nm: N=11; 60nm-80nm: N=6; >80nm: N=8. The error bars correspond to the SD. The red dashed line corresponds to a sorting ratio equal to 1. **G**: Confocal images corresponding to a GUV fusion experiment in which CHMP4B is binding in geometry (ii). Scale bar: 10µm. **H**: Quantification of the sorting ratio for 24 nanotubes from 24 GUVs in 10 independent GUV preparations and of variable diameters, corresponding to experiments of the type shown in Figure 1G. For each nanotube diameter, N measurements have been obtained: <20nm: N=11; 20nm-40nm: N=8; 40nm-60nm: N=5; 60nm-80nm: N=4; >80nm: N=11. The error bars correspond to the SD. The red dashed line corresponds to a sorting ratio equal to 1. **I**: Cryo-EM image of CHMP4B filaments polymerized outside deformable membrane nanotubes. Left: Raw image, Right: Guide for the eyes. Scale bar: 50 nm. **J**: Cryo-EM image of CHMP4B filaments polymerized onto non-deformable GlaCer tubes. Left: Raw image, Right: Guide for the eyes. Scale bar: 50 nm.

LUVs have a positive Gaussian curvature, in contrast with the negative Gaussian curvature of the membranes in biological contexts where ESCRT-III usually localizes (Supplementary Fig. S1A). In order to study the effect of membrane geometry on ESCRT assembly, we have used different assays involving membrane nanotubes and corresponding to a variety of membrane geometries (Fig. 1D).

We have recently developed a novel approach based on laser-triggered fusion ^55^ that allows ESCRT-III protein encapsulation into negatively charged GUVs ^32^. We have encapsulated fluorescently-labelled CHMP4B inside a non-charged GUV (cyan) at a concentration of about 1 μM and fused it with a GUV containing PI(4,5)P2 (magenta) (Fig. 1E). A tube was pulled beforehand from the PI(4,5)P2-containing GUV (Fig. 1D, iii), generating a physiologically-relevant membrane geometry encountered by CHMP4B. Tube diameter can be tuned by changing membrane tension with a micropipette aspirating the magenta membrane GUV. Using this setup, we observed that CHMP4B was neither enriched at the tube neck, nor in the tube. We calculated the sorting ratio, i.e. the protein enrichment in the tube as compared to the GUV, based on fluorescence ^56^ (N = 64 in total for 17 nanotubes). This ratio is larger than 1 for proteins enriched and lower than 1 for proteins depleted from membrane tubes. This quantification reveals that over the large range of tube diameters that we could explore, CHMP4B is mostly excluded from tubes with a negative mean curvature (and a null Gaussian curvature) and prefers to bind homogeneously to the flat surface of the GUV membrane (Fig. 1F). Interestingly, no preferential binding was detected even at diameters corresponding to the expected preferred curvature for Snf7 filaments (diameter between 40 nm and 60 nm) ^21, 22, 24^.

We next tested binding of CHMP4B to positively curved membranes using an assay corresponding to geometry (ii) (Fig. 1D). CHMP4B proteins were initially incubated with GUVs containing PI(4,5)P2 and afterwards, a membrane nanotube was pulled outwards (Fig. 1G). In these conditions, CHMP4B exhibited a sorting ratio of the order of 1 over the full range of accessible tube diameters (N = 39 in total for 24 nanotubes) (Fig. 1H), indicating that CHMP4B can bind to tubes with a positive curvature, although no preferential affinity for this geometry has been yet reported. Moreover, absence of CHMP4B fluorescence recovery after 6 min. in FRAP experiments performed on nanotubes suggests that CHMP4B forms stable polymers when bound to the tube in these conditions (Supplementary Fig. S4).

Finally, we used cryo-EM to study the organization of CHMP4B on tubes. Our LUV preparation was generated after a crude re-suspension of a dried lipid film, which preserves PI(4,5)P2 lipids within the lipid bilayer ^57^. This methodology generates a rather heterogeneous suspension of vesicles both in size and in geometry. We found that 15 % ± 3.4 % (N = 315 vesicles) of the vesicles were spontaneously forming tubular structures in the preparation (arrows in Supplementary Figs. S3A-B). Addition of CHMP4B does not induce any significant further tubulation of the liposomes (20 ± 14.2 %, N = 214 vesicles). We have used these tubes to study how CHMP4B filaments organize on membranes with a tubular geometry. Figure 1I shows that CHMP4B filaments bind to lipid tubes, and align along the main axis of the tubes where the curvature is minimal, forming parallel structures and inducing some helicity to the tubes. We have collected images of the tubes and performed 2D averaging and classification to enhance the contrast (Supplementary Fig. S3C). 192 sections of tubes decorated by CHMP4 were hand-picked. 5 classes were generated by 2D processing. From the averages, repetitive patterns can only be discerned parallel to the axis of the tube (See Supplementary Fig. S3C, bottom row). It is also possible to generate rigid galactocerebroside (GlaCer) nanotubes tubes supplemented with EPC 10 % (wt) and 10 % (wt) PI(4,5)P2 with a uniform external diameter of 25 nm ^58^. Similarly, CHMP4B filaments polymerize on these tubular structures and tend to be aligned along the main tube axis, although some twist is visible along the filaments (Fig. 1J).

Altogether, these results do not support previous models of a stiff CHMP4B spiral acting as a loaded spring that could induce membrane bending. Rather they suggest that CHMP4B, in the absence of the other ESCRT-III proteins, flattens membranes or assembles along the main axis of tubes where the mean curvature is null.

### CHMP2B and CHMP2A/CHMP3 selectively assemble on positively-curved membranes

We next analyzed the assembly of CHMP2B and CHMP2A/CHMP3 on membranes with specific geometries.

#### CHMP2B

First, we show by HS-AFM that CHMP2B assembles onto SLBs into ring-like structures with a diameter corresponding to the peak-to-peak distance equal to 16.4 ± 3.1 nm (N = 69) (Fig. 2A). Similar structures have been reported for CHMP2A, assembled in the absence of membranes ^23^. In contrast, the *in vivo* overexpression of CHMP2B induces the formation of rigid tubular membrane protrusions stabilized by CHMP2B helical filaments ^29^, suggesting that CHMP2B can adopt different geometries upon binding to membranes. Furthermore, CHMP2B assembles into clusters that localize at the neck of nanotubes pulled from GUVs, but is not found inside tubes ^32^. This suggests that CHMP2B has some affinity for negative Gaussian curvature, but not for negative mean curvature, in contrast with *in vivo* over-expression conditions ^29^. In order to test whether CHMP2B binds as well to positively curved membranes, we incorporated CHMP2B into GUVs and employed the I-BAR protein IRSp53 to form membrane tube invaginations on another set of GUVs ^59, 60^ corresponding to the (iv) geometry (Fig. 1D). Fusion of both GUVs (Fig. 2B, left panel) demonstrated that CHMP2B co-localizes with the positively curved membranes of internal tubular structures (Fig. 2B, right panel) (N = 7). This is a consistent with an increase of the spontaneously formed tubular structures (30.2 ± 1.6 %) observed by cryo-EM after incubation of LUVs with CHMP2B. Globally, this thus shows that CHMP2B can assemble on flat membranes and on membranes with a positive mean curvature or a negative Gaussian curvature, which, however, requires the presence of CHMP4 *in vivo* ^29^.

**Figure 2:**
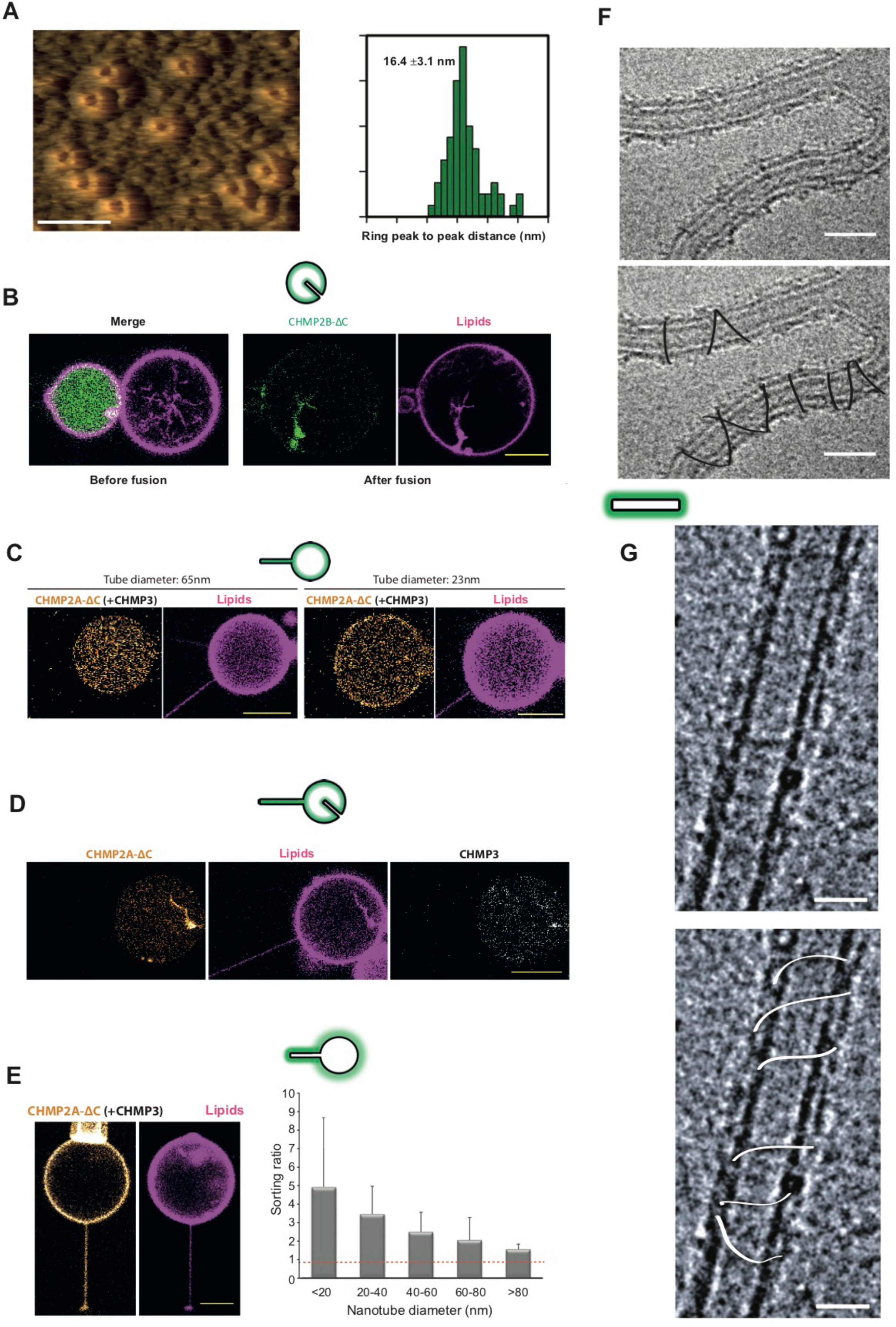
CHMP2B-ΔC assembles on positively curved tubules whereas CHMP2A-ΔC/CHMP3 binds essentially positively curved tubes and wind around them. **A**: HS-AFM image of CHMP2B-ΔC rings on a flat, non-deformable SLB. The quantification of ring diameters is shown. **B**: Confocal images corresponding to a GUV fusion experiment in which CHMP2B-ΔC is exposed to a geometry (iv) induced by the I-BAR domain of IRSp53 (non-fluorescent), tubulating the membrane when present on the exterior of the GUV. N=7. Scale bar: 10µm. **C**: Confocal images corresponding to a GUV fusion experiment in which CHMP2A-ΔC+CHMP3 are binding in geometry (iii). CHMP3 is unlabeled. Left GUV: tube diameter = 65nm. Right GUV: tube diameter = 23nm. Scale bars: 10µm. **D**: Confocal images corresponding to a GUV fusion experiment in which CHMP2A-ΔC+CHMP3 are binding in geometry (iv), showing the affinity of the assembly for internal positively curved tubes. Scale bar: 10µm. **E**: Left: Confocal images corresponding to a GUV fusion experiment in which CHMP2A-ΔC+CHMP3 are binding in geometry (ii). CHMP3 is unlabeled. Scale bar: 10µm. Right: Quantification of the sorting ratio for 24 nanotubes of variable diameters from 25 GUVs in 9 independent GUV preparations. For each nanotube diameter, N measurements have been performed: <20nm: N=20; 20nm-40nm: N=74; 40nm-60nm: N=41; 60nm-80nm: N=6; >80nm: N=4. Error bars correspond to the SD. The red dashed line corresponds to a sorting ratio equal to 1. Scale bar: 10µm. **F**: Cryo-EM image of CHMP2A-ΔC/CHMP3 filaments polymerized outside deformable membrane nanotubes. Top: Raw image, Below: Guide for the eyes. Scale bar: 50 nm. **G**: Cryo-EM image of CHMP2A-ΔC/CHMP3 filaments polymerized outside non-deformable GlaCer tubes. Top: Raw image, Bottom: Guide for the eyes. Scale bar: 50 nm.

#### CHMP2A/CHMP3

CHMP2A and CHMP3 have to be present together for efficiently binding negatively charged membranes ^23, 61^. We thus first studied whether CHMP2A/CHMP3 assembles inside nanotubes, using the geometry of the assay (iii) (Fig. 1D). In these experiments, only CHMP2A was fluorescently labeled and CHMP3 was kept unlabeled (see ^61^). Proteins were reconstituted inside GUVs at nearly physiological concentrations. No binding to the inner leaflet of the membrane nanotube was observed independently of the tube diameter (Fig. 2C, corresponding to tube diameters equal to 65 and 23 nm). We observe only a very weak binding, if anything, to the flat membrane of the GUV (Fig. 2C). This demonstrates that, surprisingly considering *in vivo* geometries, the CHMP2A/CHMP3 complex does not assemble on negatively curved membranes and has only a very weak affinity for membranes with a null-curvature.

However, in the presence of spontaneously formed internal tubules in the GUVs exhibiting a positive mean curvature (geometry (iv)) (Fig. 1D), we noticed a strong enrichment of the proteins on these structures (Fig. 2D). On the same line, we observed with cryo-EM that these proteins generate positive membrane curvature since the fraction of tubular structures is increased as compared to the control (31 ± 5 %) when CHMP2A and CHMP3 are added at 0.5 µM and 3 µM respectively, and they also produce short tubules (tens of nanometers long) on spherical liposomes (Supplementary Fig. S3D, N = 2 experiments).

In order to further confirm the preference of CHMP2A/CHMP3 for membranes with a positive mean curvature, we incubated GUVs with CHMP2A and CHMP3 and pulled a nanotube outwards (geometry (ii)). Here, CHMP2A associates in the presence of CHMP3 with the GUV membrane as well as the outer leaflet of the pulled membrane tube (Fig. 2E). By systematically quantifying the sorting ratio in this geometry, we confirmed that CHMP2A/CHMP3 binds to membrane with a positive mean curvature (N = 145 in total with 24 nanotubes) (Fig. 2E, right). Moreover, since the sorting ratio increases with tube curvature (the inverse of the tube radius) up to about 5 for tube diameters smaller than 20 nm, it demonstrates that the CHMP2A/CHMP3 complex polymerizes on membrane in a positive curvature-dependent matter, similarly to CHMP2B.

In agreement with these tube pulling experiments on GUVs, Cryo-EM visualization of the CHMP2A/CHMP3 assembly onto tubular membrane structures (geometry (i); Fig. 1D) revealed a helical rather loose polymer wrapping around membrane tubes perpendicularly to the main tube axis, both on spontaneous deformable tubes present in the LUV preparation (Fig. 2F) and on rigid GlaCer tubes (Fig. 2G). This further demonstrates the affinity of CHMP2A/CHMP3 for membranes with a positive mean curvature, which is different from the linear arrangement of CHMP4B along the main axis of tubes (Figs.1I, 1J).

### CHMP2A/CHMP3 and CHMP2B in combination with CHMP4B selectively assemble on membranes with negative Gaussian curvature and remodel membranes

#### CHMP4B/CHMP2B and CHMP4B/CHMP2A/CHMP3 complexes are recruited inside the neck of membrane nanotubes

*In vivo*, ESCRT-III complexes essentially function on membranes with a negative Gaussian curvature. We therefore co-encapsulated CHMP4B and CHMP2B (both fluorescent) as well as CHMP4B and CHMP2A/CHMP3 (with fluorescent CHMP4B and CHMP2A) at nearly physiological concentrations in GUVs composed of EPC and fused them with GUVs containing PI(4,5)P2 from which a tube was subsequently pulled (Fig. 1D, geometry iii).

Upon fusion, in some cases, no membrane binding was detected, probably due to a too low protein concentration. When binding occurs, both CHMP4B and CHMP2B bind to the inner leaflet of the GUV with a local enrichment of CHMP4B and CHMP2B at the neck of the nanotube (Fig. 3A) in 66 % of the cases (N = 12, 4 experiments). No protein was detected inside the nanotubes. When internal tubular structures were present inside the GUV (geometry iv) (4 GUVs), both proteins were found to be bound to these tubes.

**Figure 3:**
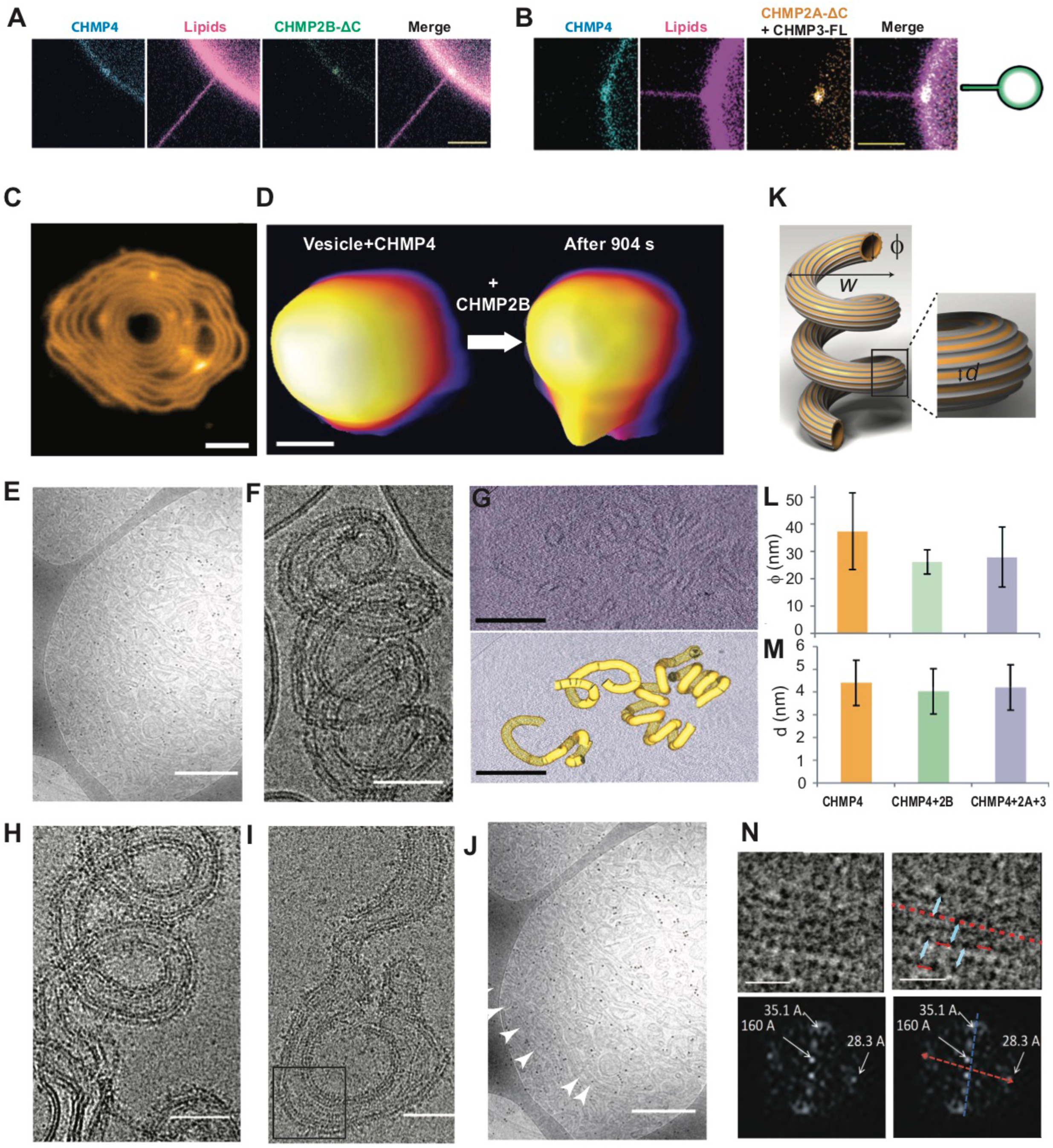
CHMP4B/CHMP2B-ΔC and CHMP4B/CHMP2A-ΔC/CHMP3 complexes are recruited inside tube neck and reshape liposomes into helical tubes. **A:** Representative confocal images of GUVs in geometry (iii) showing a preferential recruitment CHMP4B and CHMP2B-ΔC inside the tube neck. Scale bar: 5 µm. **B:** Representative confocal images of GUVs in geometry (iii) showing a preferential recruitment CHMP4B and CHMP2A-ΔC(+CHMP3) inside the tube neck. Scale bar: 5 µm. **C:** Effect of CHMP2B-ΔC addition to a CHMP4B spiral on a flat SLB, imaged by HS-AFM. Scale bar: 50 nm. **D:** Deformation of a small liposome first incubated with CHMP4B, after addition of CHMP2B-ΔC imaged by HS-AFM (corresponds to snapshots from Supplementary Movie S4). Scale bar: 100 nm. **E:** Effect of CHMP2B-ΔC addition to liposomes pre-incubated with CHMP4B, imaged by Cryo-EM at low magnification. Scale bar: 500 nm. **F:** Details of a helical tube structure induced by CHMP4+CHMP2B-ΔC. Scale bar: 50 nm. **G:** Cryo-EM tomogram of a helical tube structure formed by CHMP4+CHMP2B-ΔC (Top). Scale bar: 200 nm. Bottom: Segmentation of the tubular membrane (yellow) from the cryo tomogram displayed above. Scale bar: 200 nm. **H:** Helical tube structure induced by CHMP4+CHMP2A-ΔC+CHMP3. Scale bar: 50 nm. **I:** Helical tube structure induced by CHMP4+CHMP2A-ΔC+CHMP3. Scale bar: 50 nm. **J:** Zoom corresponding to the black frame in Fig. 3I. Scale bar: 20 nm. **K:** Scheme of the helical tubes: *w* is the width of the spiral, *ϕ* the diameter of the tube and *d* the distance between adjacent parallel filaments on the tube. **L:** Measurement from the Cryo-EM images of the tube diameters *ϕ* for CHMP4 (N=43), CHMP4/CHMP2B-ΔC (N=107) and CHMP4/CHMP2A-ΔC/CHMP3 (N=233). **M:** Measurement from the Cryo-EM images of the distance *d* between filaments parallel to tube axis for CHMP4 (N=31), CHMP4/CHMP2B-ΔC (N=66) and CHMP4/CHMP2A-ΔC/CHMP3 (N=32). **N:** Top: Class-average of helical tube sections (sub-class #3 in Supplementary Table 1) formed by CHMP4/CHMP2B-ΔC. Blue arrows indicate the distance between 2 structures parallel to the tube axis and red ones indicate the distance between 2 structures perpendicular to the tube axis (See Supplementary Fig. S7F). Scale bar: 10 nm. Bottom: Fourier-Transform (FT) with the distances corresponding to the Bragg peaks. Left: Raw data. Right: The red line represents the direction of the tube axis and the blue line to the perpendicular direction along the tube section.

In the case of CHMP4B/CHMP2A/CHMP3, slightly different results were observed. When membrane binding was observed, only CHMP4B was detected on the GUVs (N = 15, 9 experiments). In about 30 % of the cases, however, we could detect a local enrichment of both CHMP4B and CHMP2A at the neck of the nanotube (Fig. 3B). All the proteins were always excluded from the interior of the nanotubes. Eventually, CHMP4B was also often strongly bound to internal tubular structures (Supplementary Fig. S5) (90 % of 10 GUVs), with only weak enrichment, if any, of CHMP2A on these tubules.

To summarize this part of our experiments, we have found that in the absence of other ESCRT partners, these minimal complexes can be recruited to the neck of membrane tube structures exhibiting a negative Gaussian curvature. In addition, they have some affinity for membranes with a positive mean curvature.

#### CHMP4B/CHMP2A/CHMP3 and CHMP4B/CHMP2B complexes transform vesicles into pipe surfaces

The complex Vps2/Vps24 was shown to induce deformation of regular Snf7 spirals assembled on non-deformable flat SLBs ^37^. We show here with HS-AFM that CHMP2B has a very similar effect on CHMP4B spirals assembled on a SLB (Fig. 3C). The CHMP4B spirals appear to lose their regularity upon addition of CHMP2B (Supplementary Fig. S6A-E and Supplementary Movie S3). Similar observations were also obtained with cryo-EM on flattened LUVs (Supplementary Fig. S7A).

We next studied in real time by HS-AFM whether membrane reshaping occurs upon ESCRT-III protein addition. In this assay, small unilamellar vesicles (SUVs) with diameters of 60-100 nm were immobilized on a mica surface and CHMP4B was added followed by CHMP2B. In these conditions, addition of CHMP4B (2 µM) did not change the spherical shape of the SUVs over a 15 min time frame (Supplementary Fig. S6F and G). However, further addition of CHMP2B (1 µM) induced the formation of an outward protrusion from the vesicle (Fig. 3D and Supplementary Movie S4) over the same time period, showing that the combination of CHMP4B and CHMP2B can mechanically deform SUVs.

These deformations were next studied with cryo-EM over a longer period of time on a large collection of LUVs. LUVs were incubated for 1 hour with CHMP4B (0.5 to 1 µM) and upon addition of either CHMP2B (0.2 to 0.5 µM, N *=* 15 experiments) or CHMP2A (0.2 to 0.5 µM) and CHMP3 (3 µM) (N *=* 13 experiments), for one additional hour induced, extensive vesicle tubulation was observed (Fig. 3E and Supplementary Fig. S7B). Close to 100% of the LUVs were tubulated under these conditions. Strikingly, both CHMP4B/CHMP2B and CHMP4B/CHMP3A/CHMP3 remodeled the vesicles into more or less regular helical tubes, like a corkscrew (a geometrical shape called a “pipe surface”) (Fig. 3F and 3H). Helical membrane tube formation by CHMP4B/CHMP2B was further confirmed by cryoET (Fig. 3G). Both CHMP4B/CHMP2B and CHMP4B/CHMP2A/CHMP3 helical membrane tubes revealed parallel filaments, following the spiral tube axis (Figs. 3F, 3H and 3I). The sequence of protein addition is essential to trigger this type of membrane reshaping. When CHMP4B and CHMP2B, or CHMP2A/CHMP3, are added simultaneously, or when CHMP4B is added after CHMP2B (N = 2 experiments) or CHMP2A/CHMP3 (N = 2 experiments), the helical tubular deformations no longer occur and only flat spirals (Supplementary Figs. S7C, S7D and S7E) are observed, similar to the CHMP4B-only spirals (Fig. 1B). Hence, CHMP4B has to assemble first on liposomes in order to nucleate the helical membrane tube deformation by either CHMP2B or CHMP2A/CHMP3.

Figure 3K schematizes the pipe surface of the membranes upon binding of the ESCRT-III proteins, where *w* is the width of the spiral, *ϕ* the diameter of the tube and *d* the distance between adjacent parallel filaments on the tube. We found that the width *w* is conserved and equal to 115.1 ± 16.2 nm (N = 66) for CHMP4B/CHMP2B and 110.9 ± 20 nm (N = 66) for CHMP4B/CHMP2A/CHMP3. The diameters of the tubular structures are displayed in Figure 3L. The diameters of the tubes decorated by CHMP4B only (*ϕ =* 37.4 ± 14 nm, N = 43) are significantly larger than the tubes induced in combination with CHMP2B (*ϕ =* 26.2 ± 4.4 nm, N=107) or CHMP3/CHMP2A (*ϕ =* 27.9 ± 11 nm, N = 233), suggesting that the final organization of the proteins on the membrane constricts the tubes. In contrast, the distance between filamentous structures parallel to the tube axis is similar for CHMP4B/CHMP2B samples (*d* = 4.2 ± 1.1 nm (N = 60)), for CHMP4B/CHMP2A/CHMP3 samples (*d* = 4.4 ± 1.2 nm (N = 66)) and for CHMP4B only (*d* = 4.4 ± 0.8 nm (N = 31)) (Fig. 3M). In addition, striations perpendicular to the long axis of the tubes are also present (Fig. 3J arrow). To further quantify this observation, we have performed alignment and classification out of 350 sub-portions of tubes for the CHMP4B/CHMP2B sample (Fig. 3N, top). The 10 resulting classes and the resulting Fourier transformation show unambiguous evidence for periodical structures both along the axis of the tubes (red line) and perpendicular to the tube (blue line) (Supplementary Fig. S7F). In Fourier space, all of the classes generated peak densities along the direction related to the tube axis (red) (Fig. 3N, bottom and Supplementary Fig. S7F), corresponding to a repetitive pattern of consecutive filaments with a mean distance of 3.5 ± 0.3 nm, averaged from the 10 classes, similar to our image analysis (Fig. 3M). In addition, diffraction peaks indicating a repeated distance perpendicular to the axis of the tubes (blue) (Fig. 3N) were also present for 20 % of the obtained classes (3 classes) (Supplementary Table S1). We obtained 3.2 ± 0.4 nm. Note that no such orthogonal structures were detected with CHMP4B only (Supplementary Fig. S3C). Hence, two sets of perpendicular filaments are bound to the pipe surface: a first set of parallel filaments along the main axis of the helical tube and locally, on about 20% of the total tube length, a second set of perpendicular filaments, possibly influencing its diameter (Fig. 3L).

**Table 1:** Buffer used for CHMP4B fusion experiments.

| Buffer<br>Composition (mM) | PIP2 Encapsulation<br>buffer | 4B<br>encapsulation buffer | External<br>buffer |
| --- | --- | --- | --- |
| NaCl | 10 | 0 | 85 |
| Tris | 25 | 25 | 25 |
| Sucrose | 250 | 100 |  |
| Glucose |  |  | 100 |

**Table 2:** Buffer used for CHMP2A+CHMP3 fusion experiments.

| Buffer<br>Composition (mM) | PIP2 Encapsulation<br>buffer | 2A+3<br>encapsulation buffer | External<br>buffer |
| --- | --- | --- | --- |
| NaCl | 25 | 50 | 50 |
| Tris | 25 | 25 | 50 |
| Sucrose | 175 | 100 |  |
| Glucose |  |  | 100 |

Both CHMP2A and CHMP3 stock solution are at $150 \text{ mM NaCl}$ . After addition of $80 \mu\text{l}$ of CHMP2A and $50 \mu\text{l}$ of CHMP3, the encapsulation mixture has a NaCl concentration of
~90mM. After fusion with a PI(4,5)P<sub>2</sub>-containing vesicle of equal size, the NaCl concentration drops to ~45mM.

**Table 3:** Buffer used for CHMP4B+CHMP2A+CHMP3 fusion experiments.

| Buffer<br>Composition (mM) | PIP2<br>Encapsulation buffer | 4B+2A+3<br>encapsulation buffer | External<br>buffer |
| --- | --- | --- | --- |
| NaCl | 10 | 0 | 85 |
| Tris | 25 | 25 | 25 |
| Sucrose | 250 | 100 |  |
| Glucose |  |  | 100 |

**Table 4:** Buffer used for CHMP4B+CHMP2A+CHMP3 fusion experiments.

| Buffer<br>Composition (mM) | PIP2<br>Encapsulation buffer | 4B+2B<br>encapsulation buffer | External<br>buffer |
| --- | --- | --- | --- |
| NaCl | 10 | 0 | 85 |
| Tris | 25 | 25 | 25 |
| Sucrose | 250 | 100 |  |
| Glucose |  |  | 100 |

**Table 5:** Proteins concentration in fusion experiments.

| Protein or protein<br>combination | Final protein<br>concentration<br>(after fusion) |
| --- | --- |
| CHMP4B | 1µM |
| CHMP2A+CHMP3 | 1μM+1μM |
| CHMP2B | 1μM |
| CHMP4B+CHMP2A+CHMP3 | 0.8μM+0.4μM+0.4μM |
| CHMP4B+CHMP2B | 0.8μM+0.8μM |

To uncover the ultrastructure of the ESCRT-III proteins at nanometer scales, we carried out cryoET, followed by sub-tomogram averaging, of CHMP4/CHMP2B-induced tubular structures. Four different populations of filaments decorating the pipe-like architecture could be identified in the whole dataset. The first, comprising one third (34 %) of the analyzed structures, does not display any organized patterned architecture. The second (Figure 4A), is composed of individual protein filaments that decorate the pipes in a homogeneous fashion. This group was the predominant ordered ultrastructure observed in the sample (33 %). Indeed, after averaging lipid tubes using their whole sections (Supplementary Fig. S8A), we observed that 16 filaments decorate the tubular structure in a periodic and regular pattern, independent of the curvature (Supplementary Fig. S8A, Supplementary Movies S5 and S6). The reconstruction was obtained at a resolution of 28.3 Å (See FSC curve in Supplementary Fig. 8B). The third population consists of paired filaments that were found in only 3% of the dataset, (Fig. 4B) and its final average was determined at a resolution of 30.8 Å (see FSC, Supplementary Fig. S8B) (Supplementary Movie S7), Interestingly, multireference analysis (MRA), resulted in different classes grouped according to their distribution along the pipe (Supplementary Fig. S8C: class 1 and class 2). Class 1 corresponds to negatively curved portion of tubes (inner side), where filaments are scarce (Supplementary Fig. S8D). Class 2 corresponds to filaments bound to the outer side of tubes (positive curvature) with a higher density, suggesting that paired ESCRT filaments have a higher affinity for positively curved membranes. A similar asymmetric distribution has been described in a recent report on yeast ESCRTs ^62^. The fourth population, comprising the remaining 30% of the dataset, is composed of filaments bridged by protein connections perpendicular to the main axis of the tubes (see Fig. 4C, arrows) (Supplementary Movie S8). Its final average was determined at 28.3 Å resolution. This set of bridging proteins is most likely related to the striations perpendicular to the main axis of the lipid tubes visualized in Figure 3J, highlighted by diffraction spots perpendicular to the main axis of lipid tubes (Fig. 3N).

**Figure 4:**
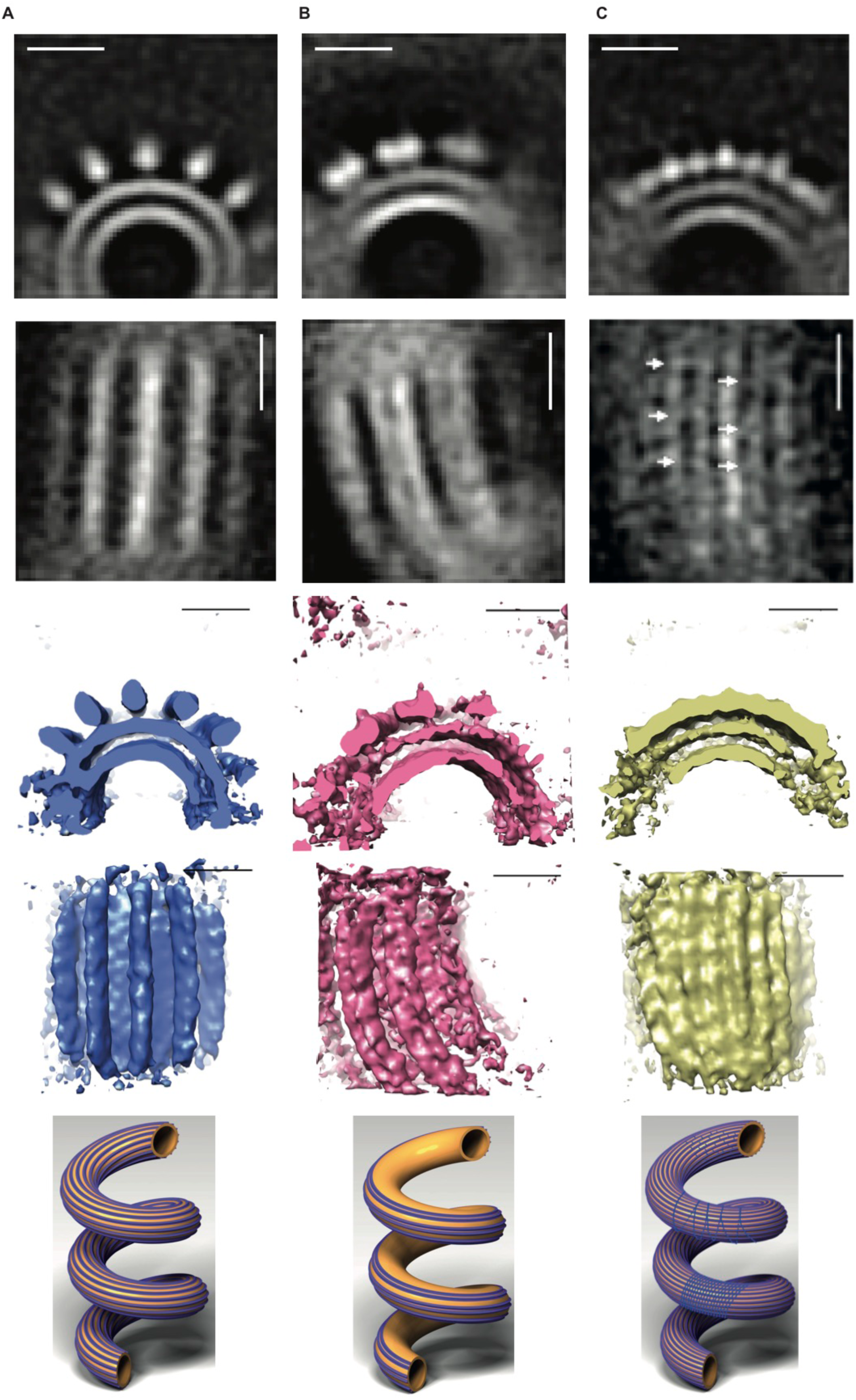
Sub-tomogram averaging of CHMP4B/CHMP2B decorated pipes. Populations of ESCRT filaments bound to tubular membranes resulting from sub-tomogram averaging. Upper line: orthoslices viewed from the cross sections of tubes. Second line: orthoslices viewed from the top of tubes. Third line: 3D reconstructions viewed from the cross sections of tubes. Fourth line: reconstructions viewed from the top of tubes. Bottom line: schematic representations of the CHMP4B/CHMP2B-decorated pipes. Scale bars: 10 nm. **A:** Single ESCRT individual filaments bound to lipid tubes. Inner tube diameter: 14.8 nm, outer tube diameter: 21.7 nm. Protein density diameter: 30.2 nm. Reconstruction from 1721 particles. **B:** Paired ESCRT filaments bound to lipid tubes. Inner tube diameter: 15.4 nm, outer tube diameter: 21.7 nm. Reconstruction from 524 particles. **C:** High density of filaments bound to lipid tubes. Inner tube diameter: 18.5 nm, outer tube diameter: 25.4 nm. Reconstruction from 381 particles. Arrows point to structures perpendicular to the tube axis.

Taken together, our analyses demonstrate that the observed macroscopic tubulation into a corkscrew-like architecture is driven by distinct nanometer ultra-structures of ESCRT filaments.

## Discussion

Membrane remodeling by ESCRT-III polymers implicates in many cases negative Gaussian membrane curvature. It has been shown that ESCRT-III assembles at or inside bud necks of endosomal vesicles ^14^, enveloped viruses ^43, 45, 48^ and within the cytokinetic midbody ^49, 52^. Less is known about membrane shape requirements or effects of ESCRT-III recruitment during the other ESCRT-catalyzed membrane remodeling processes. Interaction with positively curved membranes has been so far only reported for ESCRT-III CHMP1B that forms helical structures on membrane tubes *in vitro* and *in vivo* on endosomal tubular extensions ^63, 64^. Our results reveal that neither CHMP4B, CHM2A/CHMP3 nor CHMP2B have on their own affinity for membranes with a mean negative curvature. Instead, we show that CHMP4B assembles into spirals on flat membranes similar to yeast Snf7 ^22^. However, CHMP4B does not deform membranes on its own. Polymerization into spiral structures on the positively curved membranes of LUVs only leads to membrane flattening, but no membrane buckling occurs under these conditions in contrast to theoretical predictions ^22, 54^. Upon incorporation of CHMP4B inside GUVs with membrane tubes pulled outwards, CHMP4B does not concentrate at the tube as observed at the bud neck under CHMP2 double knockdown HIV-1 budding conditions ^43^. Thus, functional CHMP4B recruitment to membrane necks may require prior assembly of ESCRT-I and ESCRT-II complexes or Alix that coordinate the assembly of CHMP4B filaments ^65, 66^ or enhance CHMP4B affinity for the membrane, rather than CHMP4B having a preference for certain membrane curvatures ^34, 35^. In contrast, CHMP4B assembles on the outside of membrane tubes, thereby forming filaments parallel to the tube axis where mean membrane curvature is null. The large-scale twisting of the CHMP4B-decorated flexible membrane tubes further supports the helical nature of the CHMP4B filaments.

Strikingly, we have shown that both CHMP2A/CHMP3 and CHMP2B do not assemble inside tubes but rather polymerize on the outer side of membrane tubes. Moreover, high positive curvature enhances CHMP2A/CHMP3 polymerization that forms helical filaments wrapping around tubes, perpendicular to the tube axis. Although the structure of the polymers appears more lose than for CHMP1B ^30^, it thus establishes that not only CHMP1B interacts with positive curved membranes, but also ESCRT-III core members that have been implicated *in vivo* in remodeling with an opposite membrane geometry.

Only the combination of CHMP4B/CHMP2B or CHMP4B/CHMP2A/CHMP3 is found occasionally enriched at the neck of membrane nanotubes consistent with the proposal that a CHMP4 polymer forms a platform for downstream ESCRT-III assembly ^13, 43^. In addition, CHMP2A-CHMP3 and CHMP2B systematically remodel CHMP4B-bound LUVs into regular helical tubes (pipe surfaces). 3D helical polymeric structures have also been reported in the absence of membranes for the assembly of Snf7, Vps2 and Vps24 ^24, 67^, and CHMP2A/CHMP3 ^23^. For both CHMP4B/CHMP2B or CHMP4B/CHMP2A/CHMP3, the width *w* of these helices is of the order of 110 nm, thus about twice larger than the preferred one for Snf7 ^22, 24, 68^ or CHMP4B ^27, 28^. These tubular structures have a smaller diameter than the occasional tubes with CHMP4B on their surfaces, suggesting particular mechanical properties of the mixed polymers. The mechanism behind such a massive membrane remodeling is yet unclear. Using sub-tomogram averaging, we could sort the architecture of the CHMP assemblies on the pipe surfaces at nanometer scales in 2 main categories, single filaments and filaments with perpendicular connections, and a minority made of double filaments, although all leading to the same macroscopic membrane geometry. This indicates that only a limited set of filaments may be required to induce this helical tubular structure. The resolution of the current structures does not permit to conclude whether the filaments are formed by open or closed CHMP conformations ^15, 16, 30, 31^. Using budding yeast ESCRT-III proteins, a recent preprint reports that Snf7, VPS2 and Vps24 forms similar corkscrew-like membrane tube structures ^62^, suggesting that this remodeling capacity of the ESCRT-III is conserved among species. However, in this report, the global membrane shape transformation was essentially attributed to filament doublets. This type of structure represents only a very minor fraction of the organizations that we have observed with human ESCRT. In contrast, on at least one third of the surface of the spiraled tubes, we observe a combination of filaments aligned along the helical tube axis crosslinked by orthogonal structures. This sort of scaffold that combines both trends of CHMP4B to form a wide spiral organization and of CHMP2A/CHMP3 to wrap around tubes, can explain the emergence of the pipe surface geometry for the membrane, although what sets the tube diameter is unclear. In addition, a large fraction of the surface is covered with single filaments regularly distributed around the tube diameter, without any connection between them. Continuity of the surface and of the filaments may explain why the tubular geometry is maintained in the absence of connections between the spiraled filaments.

Although the helical tubular shape of the membrane may be far from membrane geometries where these core ESCRT-III proteins have been localized *in vivo*, it nevertheless reveals the mechanical stresses that these protein assemblies exert on membranes. The pipe surface represents the membrane shape that minimizes the mechanical energy of the system in the absence of any external constraints. It also shows the capacity of the proteins to assemble onto an “outside-in” geometry, similarly to CHMP1B and IST1 ^6^. Nevertheless, when overexpressed in human cells, CHMP4B induces tubules with an “in-outside” geometry ^6, 27^. *In vivo*, CHMP4B is recruited to the membrane via CHMP6, which in turn is recruited by ESCRT-II and ESCRT-I ^13, 65^ or by Alix ^69, 70^. The ESCRT-III proteins may be then forced to assemble in a non-optimal geometry and in return exert mechanical forces on the neck structure to release frustration. How these reciprocal interactions combined to the action of the VPS4 ATPase lead to membrane scission remains to be established. But, our work provides new insight on how mechanics and geometry of the membrane and of the ESCRT-III assemblies can generate forces to shape a membrane neck.

## Methods

### 1) Materials

#### Reagents

Common reagents were purchased from VWR reagents. L-α-phosphatidylcholine (EPC, 840051P), 1,2-dioleoyl-sn-glycero-3-phosphocholine (DOPC, 850375P), Cholesterol (700000P), 1,2-dioleoyl-*sn*-glycero-3-phosphoethanolamine (DOPE, 850725P), 1,2-dioleoyl-*sn-*glycero-3-phospho-L-serine (DOPS, 840035P), L-α-phosphatidylinositol-4,5-bisphosphate (PI(4,5)P2, 840046P), 1,2-dioleoyl-sn-glycero-3-phosphoethanolamine-N-(biotinyl) (PE-Biotin, 870282P) and 1,2-dioleoyl-sn-glycero-3-phosphoethanolamine-N-(lissamine rhodamine B sulfonyl) (Rhod-PE, 810150P) were purchased from Avanti polar.

Gold nanorods Streptavidin-conjugated gold nanorods (C12-10-850-TS-DIH-50) were purchased from Nanopartz™.

Streptavidin-coated polystyrene beads (diameter 3.2 μm) for the tube pulling experiments were purchased from Spherotech.

##### Recombinant proteins

CHMP3 (full length) was expressed in *Escherichia coli* BL21 cells for 3 h at 37°C ^15^. Briefly, cells were harvested by centrifugation (4000 g for 20 min at 4°C) and the bacterial pellet was resuspended in 50 ml of binding buffer A (20 mM Bicine pH 9.3, 300 mM NaCl, 5 mM imidazole, 1% CHAPS/1 mM PMSF). The bacteria were lysed by sonication and CHMP3-FL was purified by Ni^2+^ chromatography. A final gel filtration chromatography step was performed in buffer B (20 mM Hepes pH 7.6, 150 mM NaCl).

CHMP2A-ΔC was expressed as MBP-fusion protein in *Escherichia coli* BL21 cells ^19^ for 1 h at 37°C. Cells were harvested by centrifugation (4000 ***g*** for 20 min at 4°C) and the bacterial pellet was resuspended in 50 ml of binding buffer C (20 mM Hepes pH 7.6, 300 mM NaCl, 300 mM KCl). The bacteria were lysed by sonication, and CHMP2A-ΔC was purified on an amylose column. CHMP2A-ΔC was labeled overnight at 4°C with Alexa Fluor 405 NHS Ester (Thermo Fisher Scientific) using a molar ratio (Alexa Fluor:protein) of 2:1. A final gel filtration chromatography step was performed in a buffer B. CHMP3-FL and CHMP2A-ΔC were concentrated to 20 μM, and immediately frozen in liquid nitrogen with 0.1% of methyl cellulose (Sigma-Aldrich) as cryo-protectant. All aliquots were kept at −80°C prior to experiments.

CHMP2B-ΔC contains amino acids 1–154 and a C-terminal SGSC linker for cystein-specific labeling. CHMP2B-FL contains the full-length sequence with a C-terminal SGSC linker for cystein-specific labeling ^32^. Both proteins were expressed in *Escherichia coli* BL21 cells for 4 h at 37°C. Cells were lysed by sonication in buffer D (50 mM Tris-HCl pH 7.4, 1 M NaCl, 10 mM DTT and protease inhibitor (Complete EDTA free, Roche) at the concentration indicated by the manufacturer) and the soluble fraction was discarded after centrifugation (50,000 ***g***, 20 min, 4°C). The pellet was washed three times with buffer E (50 mM Tris-HCl pH 7.4, 2 M urea, 2% Triton X-100 and 2 mM β-mercaptoethanol). The last wash was performed in absence of urea and Triton X-100. The extraction of CHMP2B was performed in 50 mM Tris-HCl pH 7.4, 8 M guanidine, 2 mM β-mercaptoethanol overnight at 4°C. After centrifugation (50,000 ***g***, 20 min, 4°C), CHMP2B was purified by Ni^2+^-chromatography in buffer F (50 mM Tris-HCl pH 7.4, 8 M urea). The protein was eluted in 50 mM Tris-HCl pH 7.4, 8 M urea, 2 mM β-mercaptoethanol, 250 mM imidazole. Refolding was performed by rapid dilution of CHMP2B into buffer G (50 mM Tris-HCl pH 7.4, 200 mM NaCl, 2 mM DTT, 50 mM L-glutamate, 50 mM L-arginine) and a final concentration of 2 μM. CHMP2B was concentrated by passing it over a Ni^2+^ column in buffer H (50 mM Tris-HCl pH 7.4, 200 mM NaCl) and eluted in buffer I (50 mM Tris-HCl pH 7.4, 300 mM NaCl, 250 mM imidazole). CHMP2B was labeled overnight at 4°C with Alexa Fluor 488 C5 Maleimide (Thermo Scientific) with a molar ratio (Alexa Fluor:protein) of 2:1. A final gel filtration chromatography step was performed on a superdex75 column in buffer J (50 mM Tris-HCl pH 7.4, 100 mM NaCl). Both CHMP2B-FL and CHMP2B-ΔC were concentrated to 20 μM, and immediately frozen in liquid nitrogen with 0.1% of methyl cellulose (Sigma-Aldrich) as a cryo-protectant. All aliquots were kept at −80°C prior to experiments.

CHMP4B was expressed as MBP fusion protein ^20^ in *Escherichia coli* BL21 cells for 2 hours at 37°C. Cells were harvested by centrifugation (4000 ***g*** for 20 min at 4°C) and the bacterial pellet was resuspended in 50 ml of binding buffer K (50 mM Hepes pH 7.6, 300 mM NaCl, 300 mM KCl). The bacteria were lysed by sonication. The CHMP4B protein was purified on an amylose column. CHMP4B was labelled overnight at 4°C with Alexa 555 succimidyl ester or 633 succimidyl ester (Thermo Fisher Scientific) using a molar ratio (Alexa Fluor:protein) of 2:1. A final gel filtration chromatography step was performed in the buffer K. CHMP4B were concentrated to 15 μM and immediately frozen in liquid nitrogen with 0.1% of methyl cellulose (Sigma Aldrich) as cryo-protectant. All aliquots were kept at −80°C prior to experiments.

### 2) Electron Microscopy

#### Cryo-electron microscopy: sample preparation and imaging

A lipid mixture (70 % EPC, 10 % DOPE, 10 % DOPS, 10% PI(4,5)P2) at 1 mg.mL^−1^ was quickly dried under argon for 2 minutes and next under vacuum for 30 minutes. LUVs (Large Unilamellar Vesicles) of variable size (50-500 nm) were obtained by resuspension and vortexing of the lipid film after addition of a buffered solution to reach a final concentration of 0.1 mg.mL^−1^. Different combinations of CHMP proteins were incubated with the vesicles at room temperature for one hour. A 4 µL drop of the solution was deposited on a glow discharged lacey carbon electron microscopy grid (Ted Pella, USA). Most of the solution was blotted away from the grid to leave a thin (less than 100 nm) film of aqueous solution. The blotting was carried out on the opposite side from the liquid drop and plunge frozen in liquid ethane at −181°C using an automated freeze plunging apparatus (EMGP, Leica, Germany). The samples were kept in liquid nitrogen and imaged using three different microscopes. A Tecnai G2 (FEI, Eindhoven, Netherlands) Lab_6_ microscope operated at 200 kV and equipped with a 4kX4k CMOS camera (F416, TVIPS) was used (Institut Curie). Some of the imaging was performed as well on a 200 kV FEG microscope equipped with a direct detector (Falcon camera) (Institut Pasteur). A 300 kV FEG (Field Emission Gun) POLARA microscope (FEI, Eindhoven, Netherlands) equipped with an energy filter and a direct detector (K2 camera, Gatan) was also employed (IBS, Grenoble). The imaging was performed at a magnification of 31,000 with a pixel size of 1.21 Å using a movie mode collecting 40 successive frames for a total dose of 50 electrons per Å^2^. The different frames were subsequently aligned.

#### Cryo-electron tomography

The samples were prepared as described above. 10 nm size gold beads were added to the solution before being plunge-frozen. Tilted series were collected in low dose mode, every two degrees, using a Tecnai G2 (FEI, Eindhoven, Netherlands) microscope operated at 200 kV and equipped with a 4kX4k CMOS camera (F416, TVIPS) (Institut Curie). To preserve the information and minimize irradiations at low tilt angles, the following angular scheme was applied: from 0 to 34 degrees, then from −2 to −60 degrees and finally from 36 to 60 degrees. The dose per image was 0.8 electrons per Å^2^. The imaging was performed at a magnification of 50,000 and each image was binned twice for a final pixel size of 4.26 Å. The consecutive images were aligned using the IMOD software suite ^71^. Back projection was performed using IMOD and SIRT reconstruction was carried out using Tomo3d. The segmentation was performed manually using IMOD.

The dataset used for sub-tomogram averaging consisted of 28 tilted-series collected at the Electron Microscopy Core Facility of the European Molecular Biology Laboratory (EMBL) in Heidelberg. Image acquisition was performed on a Titan Krios microscope (FEI) operated at 300 kV using a Quantum post-column energy filter and a Gatan K2 Summit direct detector controlled by SerialEM ^72^. Tilted-series were collected using the dose-symmetric scheme (Hagen et al. 2017) in the range of ±60° and a 3° angular increment and a defocus range between −1.5 and −4.25 mm. Tilt images consisted of 13 super-resolution frames with a total dose per tilted-series of 140 e^−^Å^−2^. Alignment of tilted-images based on gold fiducials, CTF estimation (CTFPlotter) and CTF correction (CTFPhaseFlip) were achieved using the IMOD suite ^71, 73^. Tomograms were reconstructed by the weighted back-projection method applied over the CTF-corrected aligned stacks in IMOD. Bin 4X (pixel size = 5.3 Å) tomograms were reconstructed using the SIRT-like filtering method with 50 iterations to facilitate identification of membrane bilayers during catalogue annotation. Protein-induced tubes were identified and annotated in Dynamo ^74, 75^ as filaments around axis. Tube radius was determined upon aligning and averaging particles cropped from the tube axis as the center of sub-volumes of 88 pixels (46.6 nm) using as alignment mask a cylinder of 22 nm radius. Then, the center of the box was displaced to the tube’s membrane surface and the selected oversampling geometry was 16 cropping points per radius separated by 6 pixels along the tube axis. Reference free subtomogram averaging was performed on subvolumes of 34^3^ nm in Dynamo.

### 3) GUVs and Nanotubes

#### Lipid mixture preparation

Lipid stock solutions were mixed at a total concentration of 1 mg/ml in chloroform with following molar ratio: 50,7 % EPC; 10% DOPS; 10% DOPE; 15 % cholesterol; 10% PI(4,5)P2; 0,2 % DSPE-PEG2000-Biotin; 0,1 % PE–Rhodamine.

#### GUV preparation – PVA gel-assisted swelling

GUVs for tube pulling experiments were prepared with the PVA gel-assisted swelling method as previously described ^76^. Briefly, PVA gel (5% Poly(vinyl alcohol)), 50 mM Sucrose, 25 mM NaCl and 25 mM Tris-HCl, at pH 7.4) was deposited on plasma cleaned (PDC-32G, Harrick) glass coverslips (18×18 mm, VWR International, France) and dried for 50 min at 60°C. 15 µl of lipid solutions at 1 mg/ml were deposited on the PVA-coated slides and residual solvent was removed under vacuum for 20 min at room temperature. The lipid film was then rehydrated with the appropriate GUV growth buffer (see Tables 1-4) at room temperature for 45 min.

#### Protein binding to the external leaflet of GUVs for tube pulling experiments

All proteins have been incubated with GUVs in a buffer containing 25 mM Tris pH 7.4 and 50mM NaCl at a concentration of 500 nM for 30’ together with 10 nM TEV, which was sufficient to cleave at least 90 % of the MBP tags in 15’ at room temperature (not shown).

#### Protein encapsulation for fusion experiments

All proteins were co-encapsulated with purified recombinant TEV protease at a final concentration of 10 nM, which was sufficient to cleave at least 90% of the MBP tags in 15’ at room temperature (not shown). Growth and observation buffers have been adjusted to each specific protein or proteins combination in order to balance the osmotic pressure. The final protein concentration and ratios between proteins is the result the balance between a number of factors, including:

- maintaining a final NaCl concentration of ∼50mM after fusion (each protein is stored in a different storage buffer and at a different concentration)
- Maintaining a sufficient amount of sucrose inside the GUV to allow sinking in the observation chamber;
- Avoiding protein inhibition: in co-encapsulation experiments, CHMP4B binding to membrane was inhibited by CHMP2A if CHMP4B/CHMP2A ratio was raised above ∼2:1, in line with previous work showing capping activity of Vps2 towards Snf7 ^37^

#### GUV growth, washing and incubation with nanorods for tube pulling experiments

This procedure was performed as previously described ^32^ (see ^77^ for a detailed protocol). Briefly, for each experiment two types of GUVs extracted from each PVA slide were mixed with the relative external buffer matching the osmolarity and centrifuged for 10 min at 1000 g. GUVs taken from the bottom of the Eppendorf were incubated with gold nanorods 20 min at room temperature and then added to the imaging chamber.

Gold nanorods Streptavidin-conjugated gold nanorods have a peak of absorption at λ=834 nm, with a tail spanning the wavelength of the optical tweezers (λ=1064 nm). The stock solution (typical concentration 1750 ppm) was diluted 1:100 upon incubation with GUVs and again diluted 1:40 when GUVs were transferred to the observation chamber.

Fusion of GUV pairs coated with the gold nanorods is achieved by bringing the GUVs hold by 2 micropipettes into close contact with micromanipulation and by locally heating the nanorods by focusing the infrared laser on the contact through the objective.

#### Tube experiments

The tube pulling experiments and analysis have been performed as previously described in ^77^ and ^56^. For experiments probing the affinity of the proteins for positive curvature, the tube was formed by bringing briefly the GUV coated with proteins in contact with a streptavidin-coated bead trapped with the optical tweezer and moved away. For experiments involving encapsulation and fusion, the tube was pulled from the PI(4,5)P2-containing GUV prior to fusion, using a streptavidin-coated bead hold by a third micromanipulator. Fusion was then performed between the GUV pair, keeping the membrane tube in place.

The values of tube diameter (in nm) were deduced from the lipid fluorescence intensities the tube in comparison with the fluorescence in the GUV, after calibration:

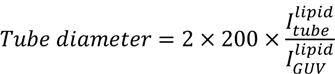

where 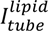 and 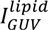 represent the fluorescence intensities of the lipids in the tube and in the GUV, respectively.

The sorting ratio S (protein enrichment in the tube) was calculated using:

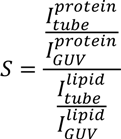

where 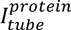 and 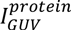 represent the fluorescence intensities of the proteins in the tube and in the GUV, respectively.

### 4) High-Speed AFM

#### Surface preparation for HS-AFM

The HS-AFM experiments on lipid bilayers were performed using SLBs composed of 60% DOPC, 30% DOPS, and 10% PI(4,5)P2. The SLBs were formed by incubating LUVs on top of freshly cleaved mica, as described in ^39^. Briefly, LUVs were thawed at room temperature and diluted to a concentration of 0.2 mg/ml in buffer (25mM Tris, pH 7.4, 50mM NaCl). Then the LUVs were incubated onto the freshly cleaved mica for 5-10 minutes, and rinsed with the same buffer afterwards.

#### HS-AFM experiments

All HS-AFM data were taken in amplitude modulation mode using a sample scanning HS-AFM [Research Institute of Biomolecule Metrology (RIBM), Japan]. Short cantilevers (USC-F1.2-k0.15, NanoWorld, Switzerland) with spring constant of 0.15 N/m, resonance frequency around 0.6 MHz, and a quality factor of ∼2 in buffer were used. The cantilever-free amplitude is 1 nm (3 nm for imaging liposomes), and the set-point amplitude for the cantilever oscillation was set around 0.8 nm (2.7 nm for liposomes). Unless mentioned, all the HS-AFM recordings were performed in buffer containing 25mM Tris pH 7.4 and 50mM NaCl.

##### HS-AFM experiments with SLBs, CHMP4B, and CHMP2B

After formation of SLBs, the surface was imaged without addition of protein. While imaging the SLB, the proteins were added to the AFM liquid chamber to reach a final concentration of 2 µM for CHMP4B, and 1 µM for CHMP2B. The formation of CHMP4B spirals on SLBs occurred within 10 minutes after incubation. To capture the effect of CHMP2B on CHMP4B spiral, the CHMP2B was only added after the formation of CHMP4B spirals was confirmed by HS-AFM imaging.

##### HS-AFM experiments with liposomes, CHMP4B, and CHMP2B

The HS-AFM experiments for dynamic membrane deformation were performed using liposomes (SUVs) composed of 50.7 % EPC; 10% DOPS; 10% DOPE; 15 % cholesterol; 10% PI(4,5)P2; 0.2 % DSPE-PEG2000-Biotin; 0.1 % PE–Rhodamine. The SUVs were obtained by sonicating a LUV mixture for 30 s. The SUVs were incubated for 5 minutes on freshly cleaved mica, and imaged under HS-AFM. Then, CHMP4B was added to reach a final concentration of 2 µM in the chamber. Later, CHMP2B (at a final concentration of 1µM) was added, but only after confirmed spiral formation (typically after 10 minutes of CHMP4B addition) on randomly formed membrane patches on mica surface. All the HS-AFM images were processed with Igor Pro with a built-in script from RIBM (Japan), and ImageJ software. Unless otherwise mentioned, all reported values are presented as mean ± SD.

## Acknowledgments

We thank Daniel Levy for support and insightful discussions, Maryam Alqabandi for her contribution on tube pulling experiments at the beginning of this work, Michael Henderson for carefully reading the manuscript. We thank Eric Nicolau for his drawing skills. This work was initiated with a grant from FINOVI (W. W., P. B.) and was supported by the ANR (ANR-14-CE09-0003-01) (W.W., P.B.), by the Institut Curie and the Centre National de la Recherche Scientifique (CNRS). W.W. acknowledges the Institute Universitaire de France (IUF) and the platforms of the Grenoble Instruct-ERIC center (ISBG; UMS 3518 CNRS-CEA-UGA-EMBL) within the Grenoble Partnership for Structural Biology (PSB). Platform access was supported by FRISBI (ANR-10-INBS-05-02) and GRAL, a project of the University Grenoble Alpes graduate school (Ecoles Universitaires de Recherche) CBH-EUR-GS (ANR-17-EURE-0003). For cryo-electron microscopy we acknowledge the support of G. Schoehn at the Grenoble Instruct-ERIC EM platform, F. Weiss and W. Hagen of the cryo-electron microscopy platform of the European Molecular Biology Laboratory (EMBL, Heidelberg) and G. Pehau-Arnaudet, M. Nilges from the UBI facility (Institut Pasteur, Paris). The Falcon II detector at the UBI facility was financed by the “Equipement d’excellence CACSICE” and the Grenoble Instruct-ERIC EM platform acknowledges support from the FRM and GIS IBiSA. Access to the EMBL cryo-electron microscopy facility was supported by iNEXT (project number 653706), funded by the Horizon 2020 program of the European Union. We further acknowledge the Cell and Tissue Imaging (PICT IBiSA, Institut Curie) platform supported by France-BioImaging (ANR10-INBS-04). N.D.F was funded by post-doctoral fellowships from the Institut Curie, the Fondation pour la Recherche Médicale and Marie Curie actions (MSCA-IF-2016 #751715 (ESCRT model)). E.M.L was supported by a post doctoral fellowship from ANR (ANR-15-CE11-0027-02). P.B. is a member of the CNRS consortium CellTiss, the Labex CelTisPhyBio (ANR-11-LABX0038) and Paris Sciences et Lettres (ANR-10-IDEX-0001-02).

## Author contributions

P.B. and W.W. conceived the study. A.B., A.C., S.Man. performed cryo-EM experiments and A.B, E.M.L. analyzed data. N.F. performed tube pulling experiments and analyzed data. W.R. supervised HS-AFM work and S.Mai performed HS-AFM experiments and analyzed data. N.M. purified and labeled CHMP proteins. A.B., N.F., S.Mai., W.R., S.Man., W.W. and P.B. wrote the paper.

## Supplementary Figure Legends

**Supplementary Figure S1: Curvatures corresponding to different membrane geometries relevant for ESCRT binding.**

The bud necks where ESCRT-III generally assemble are surfaces generally described by 2 principal curvatures that have opposite signs, thus a negative Gaussian curvature 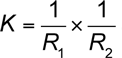. A neck shape can be close to a catenoid surface (A) when the mean curvature 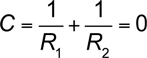, but during the fission processes, *C* and *K* can vary. For a comparison, in a tubular structure, the curvature along the axis is null (C_1_=0 or equivalently R_1_=∞), thus *K*=0, whereas in the perpendicular direction, the curvature is positive outside of the tube (thus the mean curvature *C*>0) and negative inside (*C*<0) (B).

**Supplementary Figure S2: Characterization of the CHMP4B filaments on supported lipid bilayer with HS-AFM.**

**A**: A typical example of an HS-AFM image of a CHMP4B spiral on a supported lipid bilayer.

**B**: Cross section along the color-coded lines in panel (A).

**C** and **D**: Histograms of the measured distances as indicated (as an example) by black lines in panel (B). In (**C**): distribution of the inter-filament peak to peak distances (N=134). In (**D**): distribution of central diameters of the CHMP4B spirals (N=18). The mean values ± SD are provided.

**Supplementary Figure S3: Cryo-EM control experiments in the absence of proteins and in the presence of CHMP4B or CHMP2A-ΔC/CHMP3.**

**A** and **B**: LUVs resuspended from a lipid dry film and imaged by Cryo-EM at low (**A**) and high (**B**) magnification. Scale bars: A, 250 nm; B, 50 nm. The vesicles are spherical, and approximately 15% of the samples display tubes (white arrows).

**C**: 2D analysis of lipid tubes in the presence of CHMP4B. Scale bars: 20 nm. Upper row: Boxed tube sections picked from the raw images. Middle row: Class averages generated by 2D alignment and classification. Lower row: Fourier transform corresponding to the classes above displaying diffraction peaks, characteristic of the inter-filament distance within the tube. The red dash line corresponds to the tube axis.

**D**: Cryo-EM image of LUVs incubated with CHMP2A-ΔC+CHMP3 at 0.5 µM and 3 µM, respectively. Scale bar: 100 nm. CHMP2A-ΔC/CHMP3 induces short tubules (white arrows) extending out of the vesicle.

**Supplementary Figure S4: CHMP4B polymerizes on tubes.**

No fluorescence recovery of CHMP4B is detected 6 min after photobleaching, indicating the formation of stable polymers onto the tube. Scale bar: 20µm.

**Supplementary Figure S5: CHMP4B/CHMP2A-ΔC/CHMP3 binds to positively curved membranes.**

Representative confocal images of GUVs in geometry (iv) in the presence of CHMP4B and CHMP2A-ΔC+CHMP3 showing a preferential recruitment of CHMP4B onto the internal tube. Scale bar: 20µm.

**Figure Supplementary S6: Effect of CHMP4B on liposomes and of CHMP2B-ΔC on CHMP4B spirals, studied by HS-AFM.**

**A**: A typical example of an HS-AFM image of a CHMP4B spiral deformed by the addition of CHMP2B-ΔC.

**B:** Cross-sections along the different colored lines in (A).

**C:** Distribution of inter-filament distances at the site of deformations (N=65), corresponding to the green line in (B).

**D**: Distribution of inter-filament distances at a non-deformed filament (N=51), corresponding to the red line in (B).

**E**: Distribution of the central diameters of CHMP4B spirals deformed by CHMP2B (N=22), corresponding to the blue line in (B).

The mean values ± SD are provided.

**F:** Snapshots of a small liposome in the absence of proteins, imaged by HS-AFM. Scale bar: 100 nm.

**G:** Snapshots of a small LUV incubated with 2 μM CHMP4B, imaged by HS-AFM. Scale bar: 100 nm.

**Supplementary Figure S7: Liposome shape changes induced by combinations of CHMP4B, CHMP2B-ΔC and CHMP2A/CHMP3, studied by Cryo-EM.**

**A**: Cryo-EM images of LUVs incubated with CHMP4B and then CHMP2B-ΔC. The spiral of CHMP4 is deformed by the presence of CHMP2B-ΔC. Scale bar: 100 nm.

**B**: Low magnification cryo-EM images of LUVs incubated with CHMP4B followed by CHMP2A-ΔC and CHMP3. Scale bar: 500 nm.

**C-E**: Cryo-EM images of LUVs incubated with CHMP4B and CHMP2B-ΔC added simultaneously (**C**), CHMP2B-ΔC first and then CHMP4 (**D**) and CHMP2A+CHMP3 and then CHMP4B (**E**). Scale Bars: 100 nm.

**F**: Scheme of the repeat distance along the tube diameter (red arrows) and perpendicular to the tube diameter (blue arrows). Illustration of the axes shown in the FT in Figure 3N.

**Supplementary Figure S8: Subtomogram averaging of CHMP4B/CHMP2B-ΔC on pipe surfaces.**

**A:** Average from boxing out the whole section of a tube using the data corresponding to single filaments (Figure 6, right).

**B**: Fourier shell correlation for the three reconstructions displayed in Figure 6. The resolution was determined from FSC=0.5.

**C**: Classes (1: red and 2: blue) from MRA classification using the data corresponding to paired filaments. Top: top views of tubes, Bottom: side views of tubes.

**D**: Spatial distribution on a tube of class 1 (red) and class 2 (blue) resulting from MRA classification. Class 1 particles localize onto negative curvatures while class 2 particles localize onto positively curved portion of the tubes.

Scale bars: 10 nm.

## Supplementary Table Legends

**Supplementary Table S1: Summary of the repeat distances on helical tubes obtained from cryoET class averaging and Fourier Transform.**

Distances between parallel filaments along the tube axis and repeat distances between structures perpendicular to the axis are provided for the 3 classes exhibiting regular structures in the 2 perpendicular directions.

## Supplementary Movie Legends

**Supplementary Movie S1. CHMP4B spiral oligomerized on a flat SLB, imaged by HS-AFM.** HS-AFM imaging of a CHMP4B spiral at a 1 frame/s rate. No observable change in the spiral topography was observed during imaging.

**Supplementary Movie S2. CHMP4B bound to a LUV, analyzed by cryoET.** Successive orthoslices within a typical cryo-tomogram are visualized. In the segmentation, the membranes are displayed in yellow, free CHMP4B filaments are segmented in blue, while bound CHMP4B spirals are segmented in red. Scale bar: 200 nm.

**Supplementary Movie S3. Effect of CHMP2B-ΔC on a CHMP4B spiral, as captured by HS-AFM imaging.** HS-AFM imaging at 1 frame/s of a preformed CHMP4B spiral in the presence of 1 µM CHMP2B-ΔC. It is observable that the spiral loses its structural regularity upon interaction with CHMP2B-ΔC.

**Supplementary Movie S4. Combined effect of CHMP4B and CHMP2B-ΔC on the shape of a SUV, as captured by HS-AFM.** The small liposome was imaged after incubation for 10 minutes with 1 µM CHMP2B-ΔC. Initially, there was no observable change in the physical dimension of the liposome. However, a deformation was observed after application of 1µM CHMP2B-ΔC. The images were captured at a 1 frame/s rate.

**Supplementary Movie S5**. Single ESCRT individual filaments bound to lipid tubes. Inner tube diameter: 14.8 nm, outer tube diameter: 21.7 nm. Protein density diameter: 30.2 nm. Reconstruction from 1721 particles. Frame width: 34 nm.

**Supplementary Movie S6**. Single ESCRT individual filaments bound to lipid tubes visualized onto the whole section of a tube. Frame width: 46.6 nm.

**Supplementary Movie S7**. Paired ESCRT filaments bound to lipid tubes. Inner tube diameter: 15.4 nm, outer tube diameter: 21.7 nm. Reconstruction from 524 particles. Frame width: 34 nm.

**Supplementary Movie S8**. High density of filaments bound to lipid tubes. Inner tube diameter: 18.5 nm, outer tube diameter: 25.4 nm. Reconstruction from 381 particles. Frame width: 34 nm.

